# BoxCar Assisted MS Fragmentation (BAMF)

**DOI:** 10.1101/860858

**Authors:** Conor Jenkins, Ben Orsburn

## Abstract

A recent study described the utilization of complex quadrupole isolation schemes to reduce high abundance ion saturation in Orbitrap systems. The BoxCar technique effectively democratizes MS1 scans by restricting high abundance ions from consuming as much space in the trap. This restriction allows lower abundance ions more acquisition time and can increase the signal to noise by a full order of magnitude. While effective at the MS1 level, BoxCar does not show an improvement in MS/MS acquisition as ions selected for fragmentation must come from an additional MS1 full scan in the method. In this study we describe BoxCar Assisted MS Fragmentation (BAMF), wherein ions for fragmentation are selected directly from the BoxCar scans. When utilizing BAMF, we observe the identification of ions by MS/MS that are not at all detectable in the MS1 scans of identical concentrations of peptides analyzed by standard data dependent acquisition experiments.

**Abstract Graphic:** 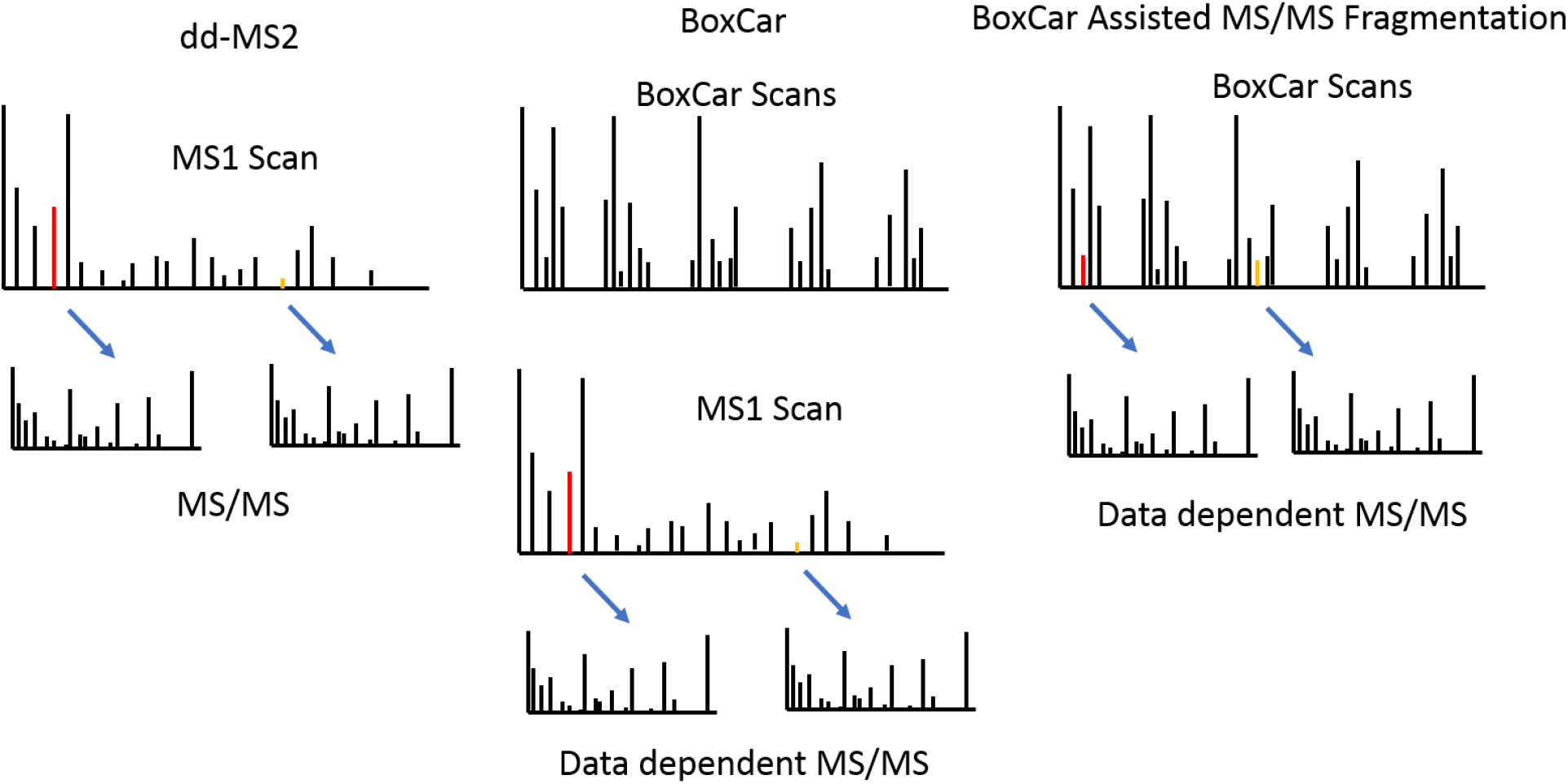

## Introduction

A recent study demonstrated the power of using advanced instrument control software for staggered multiplexed acquisition of ions for label free quantification. The method, which the authors termed BoxCar was shown to increase the dynamic range of MS1 ions to nearly 10-fold beyond traditional MS1 based data dependent shotgun proteomics.^1^ In order to use BoxCar for Q Exactive systems the operator must utilize the newly-released MaxQuant.Live software which operates in tandem with the vendor software package, Xcalibur.^2^ The newly released Exploris 480 system does, however, have full integrated BoxCar support (data not shown). To the best of our knowledge, the only way to quantitatively process BoxCar data is through the most recent releases of the MaxQuant software package which has been supplemented with an equally innovative method for normalizing intensity between BoxCar scans.

Three main drawbacks currently exist in the BoxCar methodology. The first is that it can only be utilized on the most recent quadrupole Orbitrap instruments, the Q Exactive HF and HF-X systems. In addition, in the BoxCar method only the MS1 scans benefit from the increase in dynamic range. The authors demonstrate a remarkable power in BoxCar in making peptide identification based solely on MS1 spectral libraries. The sensitivity and selectivity of MS1 based identification and quantification is well established for both the analysis of specific PTMs in complex matrices and in peptide quantification.^3^ Independent attempts to quantify resolution versus specificity confirmed that MS1 scans above a resolution of approximately 60,000 at m/z 200 can provide the same, or superior specificity as triple quad based SRM assays.^4,5^ However, the majority of modern shotgun proteomics relies on MS/MS scans to confirm peptide identification and many practitioners may not consider MS1 information sufficient for peptide identification in their experiments. In BoxCar, the selection of MS/MS spectra is performed from an additional full scan MS1 event that occurs in addition to the 3 separate BoxCar scans. While MS1 scan data benefits from the BoxCar methodology, this benefit is not passed on to the ions selected for fragmentation. A final, and perhaps most challenging drawback of the BoxCar method is cycle time. With 3 BoxCar scans and a full scan required for parent ion selection as demonstrated in the original study, the amount of time available for acquiring MS/MS spectra is reduced markedly from standard dd-MS2 experiments.

In this study we describe the design and implementation of instrument parameters inspired by BoxCar using unmodified vendor control software. While some characteristics of the BoxCar method cannot be completely replicated, we find that many of the advantages of the method can be obtained by mimicking parameters on unmodified Tribrid LC-MS systems. Furthermore, we demonstrate the capability to select ions for MS/MS fragmentation directly from the BoxCar scans and thus improving on the original technique. When utilized on a hybrid quadrupole ion trap Orbitrap system such as a Fusion or Fusion Lumos, the ability to simultaneously acquire BoxCar scans and MS/MS ion the ion trap can alleviate much of the cycle time deficit of the original experiment. All methods described in this study are posted on www.lcmsmethods.org and are freely available.

## Materials and Methods

### Samples

All experiments were carried out with the HeLa digest and Pierce Retention Time Calibration Standard (PRTC), both from Thermo Fisher. The PRTC was diluted with 0.1% formic acid to a total concentration of 50 fmol/μL. This solution of PRTC was used to reconstitute and dilute the HeLa digest standard. The HeLa standard was diluted to allow multiple injections ranging from 20ng to 500ng of peptide digest on column.

### Orbitrap Fusion Methods

All methods were written on an Orbitrap Fusion 1 system equipped with Tune version 3.0 installed in January 2018. An EasyNLC 1200 system and EasySpray source was used for all experiments with 0.1% formic acid as Buffer A and 80% acetonitrile with 0.1% formic acid as buffer B. EasySpray 25cm columns with PepMap C-18 with 2um particle size and equivalent 2cm precolumn trap was used for all experiments described. All gradients were identical for the experiment, beginning at 5% buffer B and ramping to 26% B in 51 minutes and increasing to 38% B by 81 minutes, all at 300nL/min. A steep 6-minute ramp to 98% buffer at 500nL/min was used before returning to baseline conditions. Baseline equilibration was performed automatically by the software at the beginning of each run.

All diluted samples were analyzed with the vendor’s “Universal Method” with one substitution. HCD fragmentation replaced CID to increase overall MS/MS acquisition rate^6,7^. Variations on the BAMF method are described in the results and conclusions section.

### Database searching and interpretation

Proteome Discoverer 2.2 (Thermo) was used for MS/MS analysis using a basic search workflow. Unless otherwise noted, all Orbitrap MS/MS files were searched with SequestHT using a 10ppm MS1 and a 0.02 Da MS/MS tolerance. All ion trap MS/MS spectra were searched with the same precursor tolerance and a 0.6 Da MS/MS. For the canonical human database, UniProt/SwissProt downloaded in February 2018 was parsed within the vendor software on “sapiens”. The cRAP database (https://www.thegpm.org/crap/) was used to flag contaminants in both the processing and consensus workflows.

FDR was estimated by the various nodes, Percolator, Target Decoy and Fixed Value PSM validator according to manufacturer defaults and are used as described in the results and conclusions section.

MaxQuant 1.6.3.3 was utilized for comparison of the BAMF and ddMS2 files acquired for this study as well as the BoxCar files from the original study. BAMF files will not be recognized currently as BoxCar within MaxQuant but will be recognized as “Standard” data dependent files. Utilization of the HeLa library and a secondary MCF-7 library from (Ref here) was performed as described previously.

## Results

### Designing Parallelized BoxCar Assisted Experiments

The BoxCar experiment uses the MaxQuant.Live software package and is currently only compatible with the Q Exactive HF and HF-X systems through an unlocked interface. We attempted to replicate the BoxCar methods using the Orbitrap Fusion 3.0 software vendor software. The Fusion 3.0 software has a multi-layered design with standard features shown as defaults under the subheading “Favorites”. By opening the menu options by “Show All” the user is presented with more flowchart options. If the experiment requires more options than can be completed in one logical flow chart, additional “Experiments” may be added. Control of multiple experiments is performed by managing the features “cycle time” and “loop count” within each experiment flowchart. When the logical flowchart is completed in real time by the instrument it moves to the next experiment flowchart. When all experiments are completed, the instrument returns to the first experiment.

For example, to most accurately replicate the BoxCar experiment, four experiments must be designed. These consist of three BoxCar experiments, followed by a single standard data dependent experiment flowchart. In order to simulate the BoxCars, multiplexed targeted SIM scans (msx-tSIM) can be employed and controlled independently within each experiment. However, the Fusion instrument flowchart allows data dependent scans to be performed directly from the msx-tSIM scans, as shown in Figure 1, improving both cycle time and allowing BoxCar benefits to be applied to ions selected for fragmentation. In order to clarify where our experiment differs from BoxCar, we will refer to it as BoxCar Assisted MS/MS Fragmentation (BAMF).

**Figure 1.**
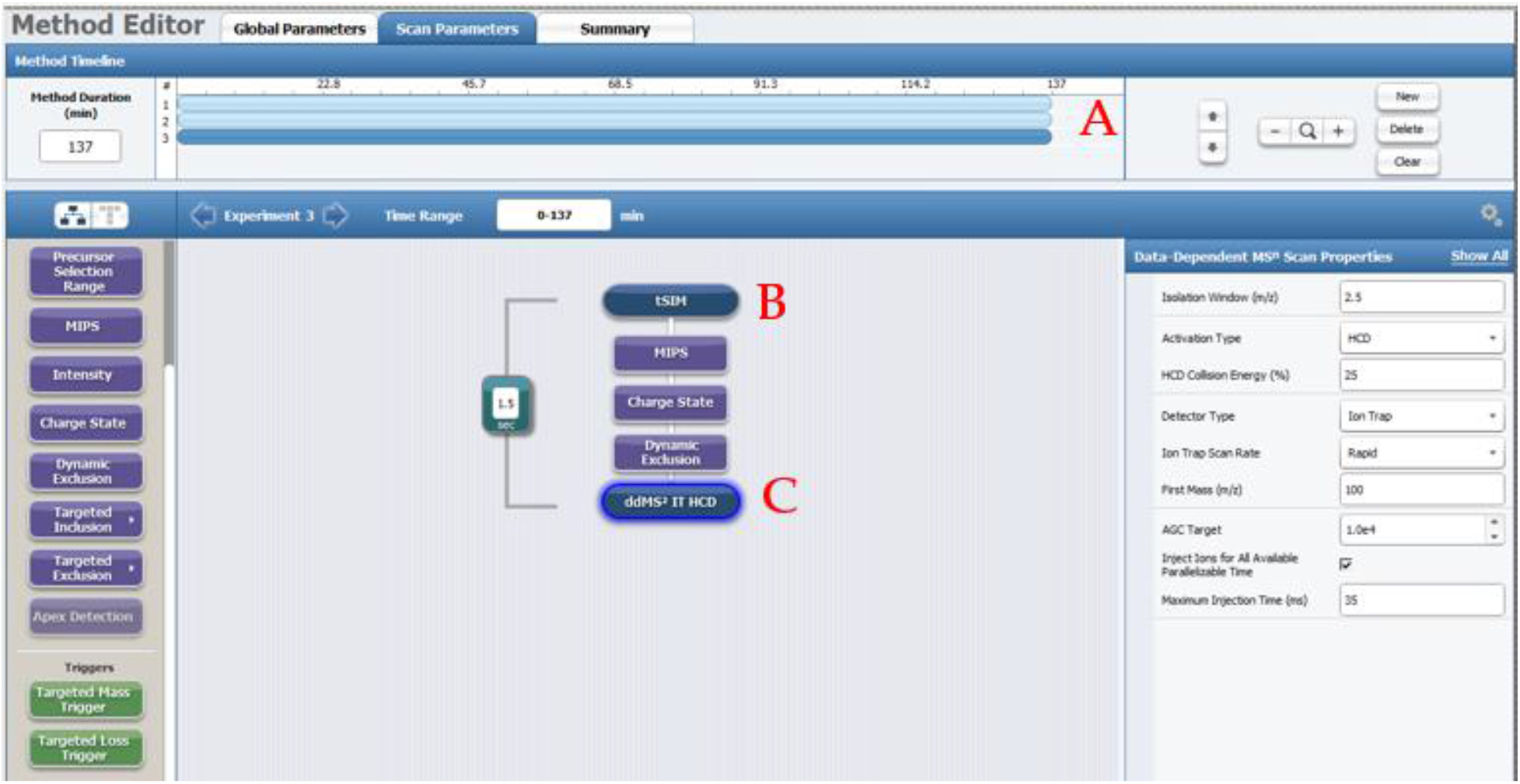
**A**. Writing the BAMF Method in Fusion 3.0 requires the use of 3 separate experiments utilizing nearly identical logical flowcharts **B**. BoxCars are written as msx-tSIM experiments multiplexed up to 10 times where the BoxCar windows are defined for each separate experiment. **C**. In order to direct MS/MS events directly from the BoxCar scans, ddMS2 is directly linked within each experiment.

Since the vendor software does not currently allow the acquisition of more than 10 msx-tSIM scans per experiment, we must use larger windows than BoxCar. When 3 experiments are utilized, 10 overlapping SIM windows were described in each experiment. The Q Exactive HF on which BoxCar was originally designed contains segmented hyperbolic quadrupoles that allow a more symmetrical isolation than traditional quadrupoles^8^. The Orbitrap Fusion 1 system contains non-segmented quads and, as such requires greater overlap between scans to minimize the loss of signal at the edges of the isolation window (data not shown).

The T-SIM windows are available in Supplemental Data 1. The full experimental methods are available in text format as Supplemental Tables 2 and 3. The original methods files are available at www.LCMSmethods.org. Figure 2 shows data produced from 3 overlapping BAMF windows demonstrating that data output is similar to the original method. As in the original technique, we experience similar democratization of the window, with no apparent observation of single ions consuming the full MS1 AGC target across all mass ranges. In addition, we find near complete utilization of the fill time to match transient is possible with BAMF, as in BoxCar, with the similar improvements in the signal to noise ratio of low abundance ions.

**Figure 2.**
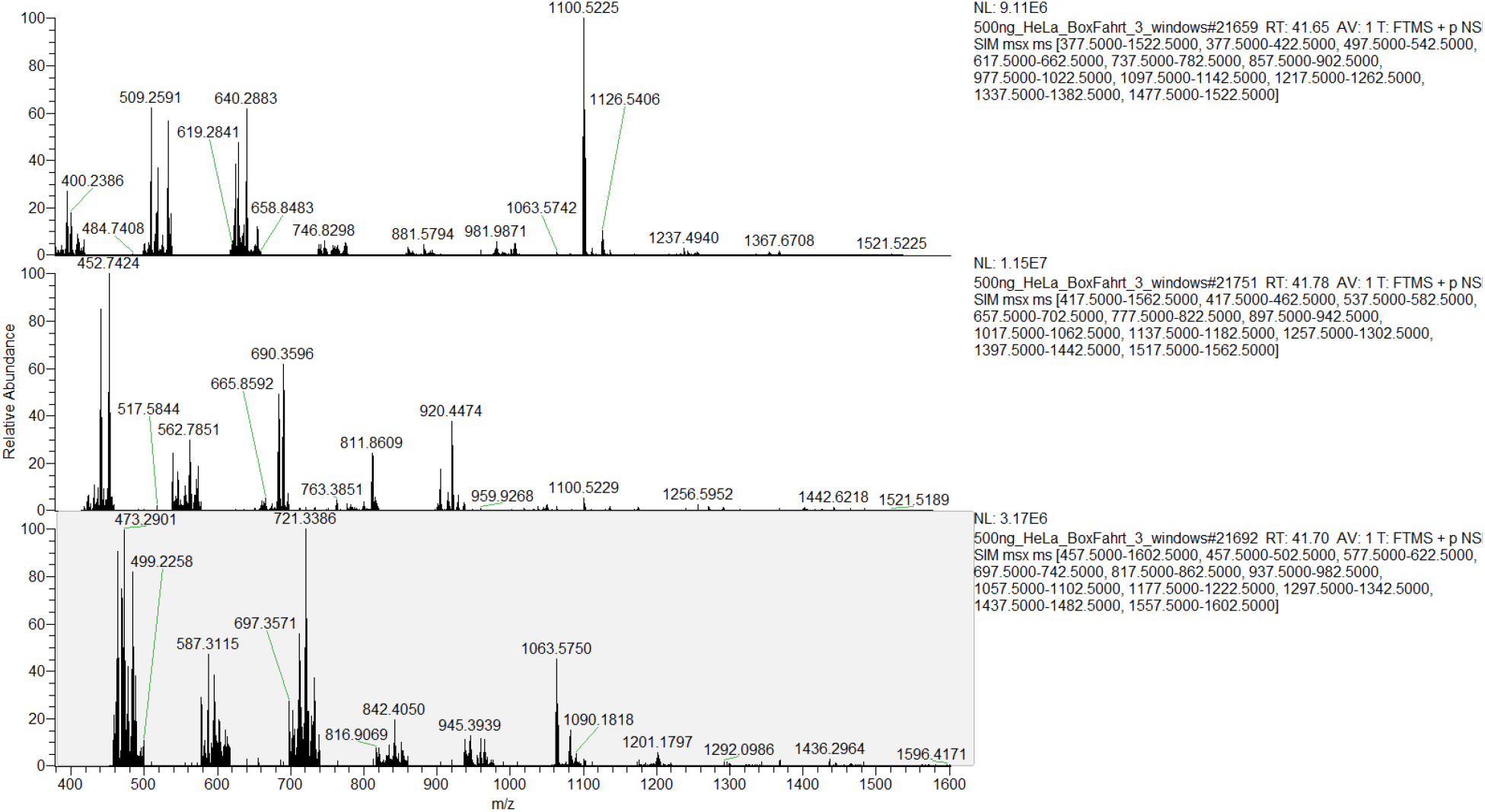
The three spectra from 3 BoxCar scans removed at subsequent time points from 500ng_HeLa_BAMF_3 demonstrating the ability of the Fusion software to acquire BoxCar data.

**Figure 2.**
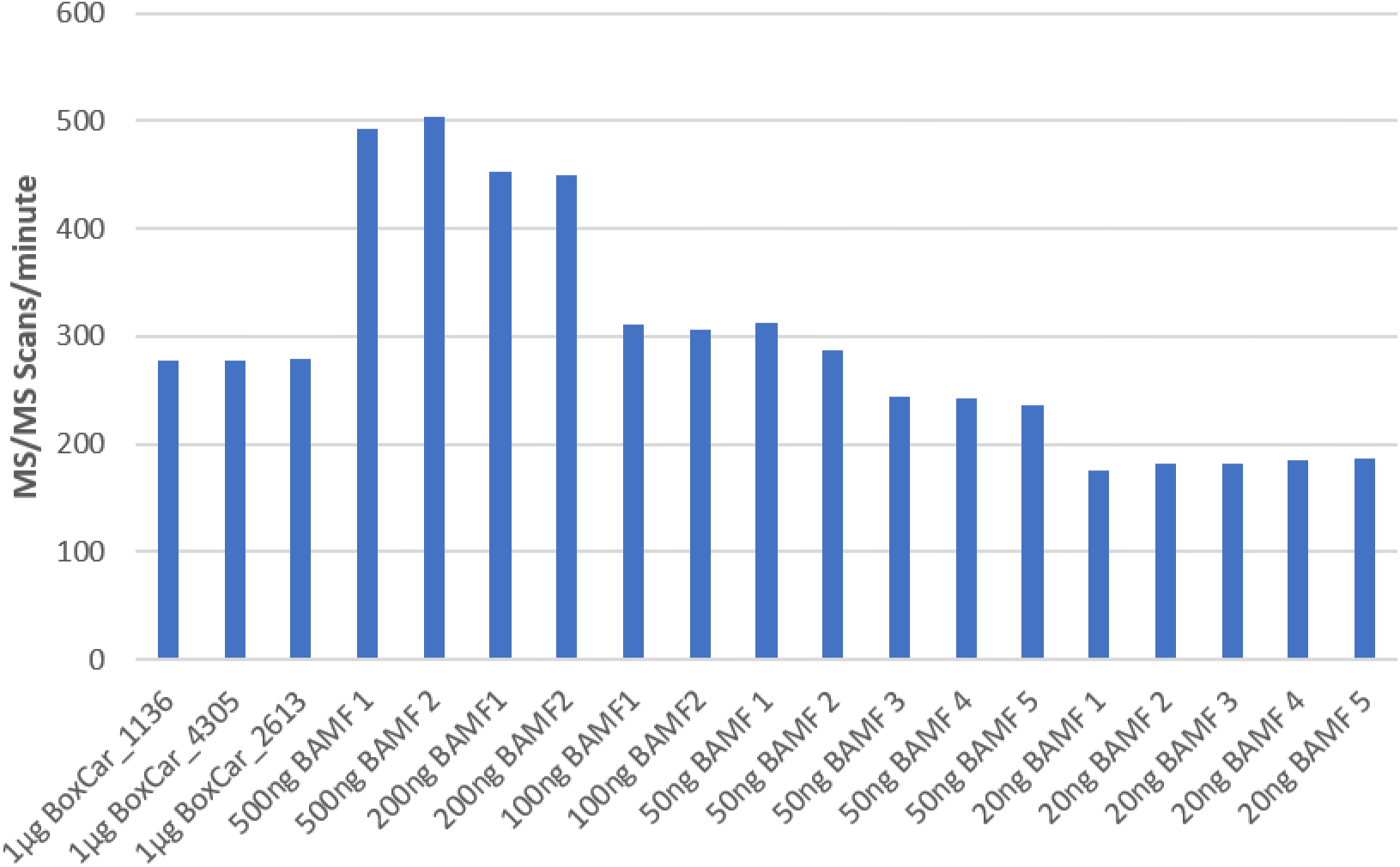
The number of MS/MS scans acquired per minute in 3 files from the original BoxCar study (left) and compared to a various on-column loads of HeLa peptides acquired with BAMF.

The Orbitrap Fusion Tribrid systems have the capability to obtain both MS1 and MS/MS spectra simultaneously by utilizing the ion trap and Orbitrap in tandem^9^. To obtain the highest possible sequencing speed, the Ion Routing Quadrupole can be used for both parent ion isolation and HCD fragmentation^7^. Figure 2 demonstrates the number of fragmentation scans obtained per minute in a dilution series of HeLa compared to the original BoxCar files. When HCD fragmentation and ion trap MS/MS scans are used, more fragmentation scans are obtained per unit time with even one tenth the total peptide load. As the concentration of peptides on column decreases, the injection time required to reach the AGC target subsequently increases. The parallelization of the two analyzers, relative increase in speed and sensitivity of the ion trap, and the lack of the full scan MS1 in BAMF offsets this in relation to BoxCar. The effect is most pronounced when relatively higher loads allows ion trap acquisition rate to be the limiting factor rather than ion injection time.

### Signal to noise improvement from BoxCar and BAMF

In the BoxCar study, the authors demonstrate an increase in signal to noise ratio of 10-fold in complex matrices. To determine whether this observation held in our methods we extracted monoisotopic ions in the same manner. Without the ability to directly quantify BAMF data automatically through the MaxQuant software we developed an alternative strategy based on data available in the literature^10^. In this study, an Intensity Based Absolute Quantification (iBAQ)^11^ analysis of the HeLa cell line generated the relative abundance of the near-complete proteome of this cell line. Using this as a metric for abundance we could compare the relative sensitivity of our dd-MS2 and BAMF experiments. The maximum tryptic digest iBAQ is Cofilin-1 with a signal of 2.57e9. In a crude metric of relative protein abundance, we distinguish proteins in this group by orders of magnitude below this intensity. For example, 4964 proteins possessed numerical iBAQ below 2.57e6, thus three orders of magnitude below maximum. As shown in Figure 3, with 20ng of HeLa digest using BAMF, 112 proteins below this mark were identified by MS/MS. This number is higher than the 70 proteins found in a file from the original BoxCar study (file 11753) despite the fact 50 times more peptide was loaded on column in the BoxCar experiment. However, equivalent protein recovery was found with 15 proteins matching the lowest protein abundances at 4 orders below maximum iBAQ.

**Figure 3.**
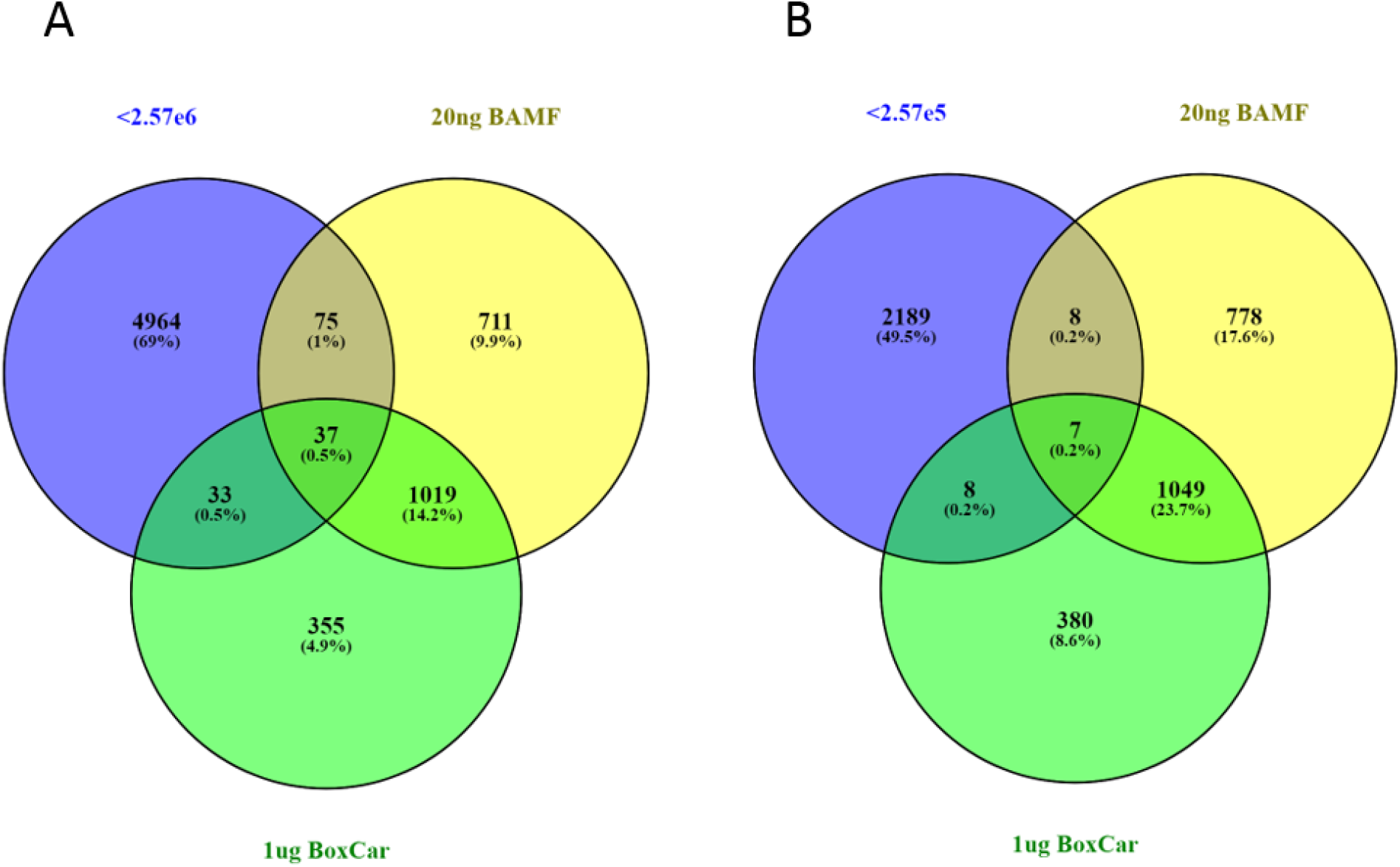
**A)** Proteins identified by MS/MS fragmentation in the 20ng BAMF run 3 compared to a file from BoxCar curated with an iBAQ 3 orders of magnitude below maximum. **B)** As in A, but 4 orders of magnitude below maximum iBAQ in this cell line.

Remarkably, we have found it possible to identify peptides fragmented with BAMF that cannot be extracted from the MS1 of our matched dd-MS2 experiments. Figure 4 is an example of one such case found by examining high scoring peptides present in BAMF, but not identified in dd-MS2. The peptide was identified from MS/MS using a 20ng injection of HeLa lysate. It can be visualized in 3 window BAMF files from 20ng through 500ng, but the XIC of even 500ng of HeLa lysate does not produce a signal for this peptide in the standard data dependent experiment. In the fragmentation of this peptide from 20ng BAMF file, 248ms of fill time is necessary to reach the ion trap AGC target further supporting the low abundance of this peptide. Cross-referencing with the HeLa iBAQ database, we find this protein at 2.3e7, more than two orders below the maximum signal.

**Figure 4.**
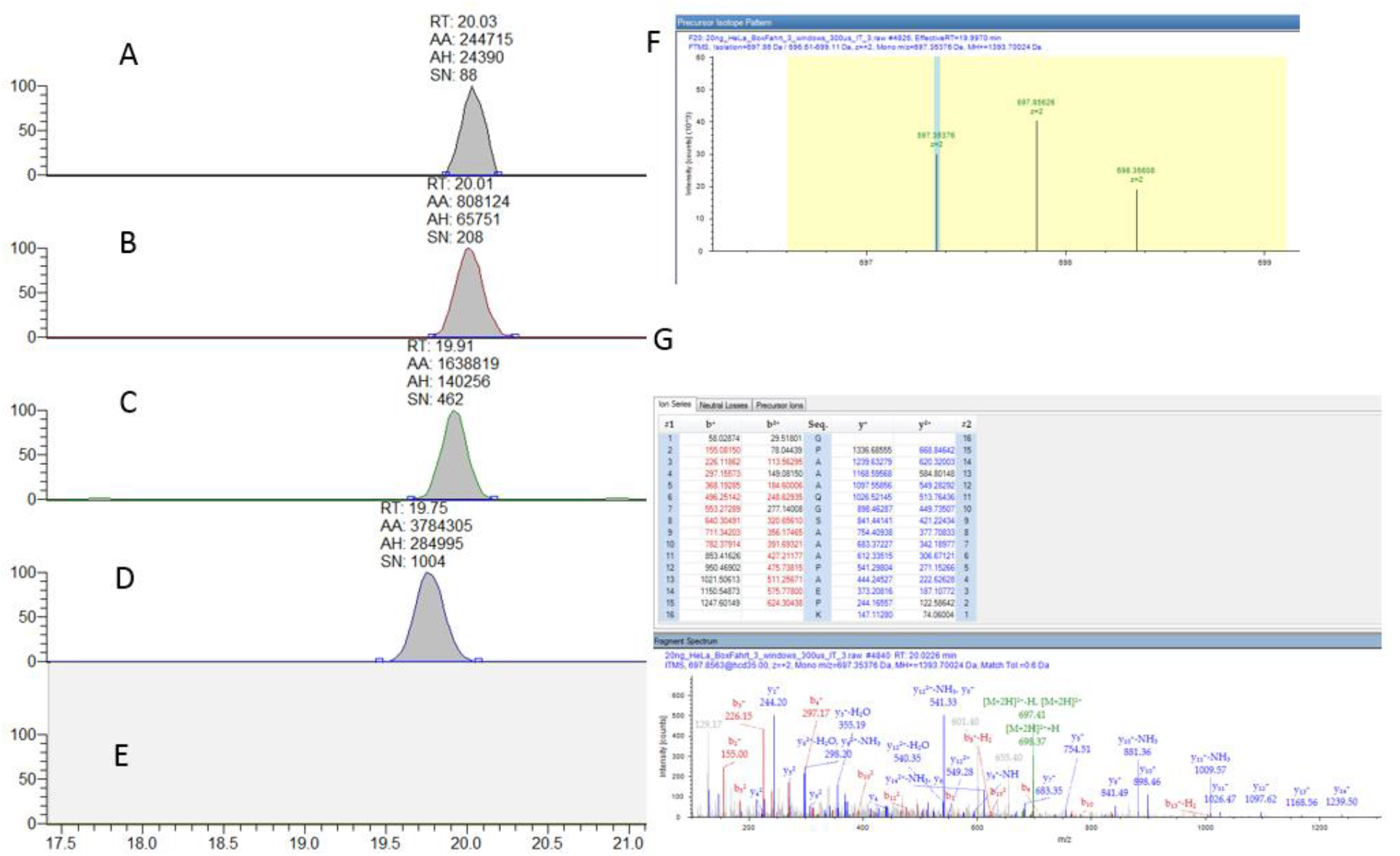
Demonstration of a peptide detected and fragmented by BAMF that cannot be extracted from dd-MS2 XIC. **A-D)** The XIC from the BoxCar window for the peptide demonstrating a near linear increase in S/N and peak area subsequent to injection volume A) 20ng B) 50ng C) 100ng D) 250ng. E) Representative XIC from the dd-MS2 experiments, 500ng dd-MS2_experiment 2 shown. **F)** The Precursor Isotope Pattern extracted for the peptide selected for fragmentation from the 20ng BAMF file demonstrates a calculated coisolation interference of 0%. **G)** The MS/MS pattern from this peptide demonstrating complete sequence coverage of this peptide.

### Peptide and Protein Identifications with BAMF

The BAMF files and BoxCar public access files were processed with both Proteome Discoverer (PD) and MaxQuant. PD was not capable of quantification and returned errors for both sets of files when using the label free quantification nodes Minora and apQuant^12^. PD can identify ions from MS/MS scans from both files using SequestHT and returned results manual verification of PSMs confirmed as appropriate. The MaxQuant version utilized could process the BoxCar files but does not currently recognizes BAMF files as BoxCar and returns errors in our hands. We selected 2 BoxCar and 2 BAMF files from each experiment and processed with the highly fractionated HeLa library generated in the original BoxCar study. Figure 5 is a summary of these results. While BAMF performed well at every dilution when MS/MS scans were matched against the FASTA file, MS1 matching was found to be inferior to the BoxCar files. While BoxCar has the advantages of dynamic isolation window control that aids in signal to noise improvements, manual examination of the BoxCar and BAMF windows suggest similar signal to noise improvements and we believe that this decrease is more reflective of file incompatibility rather than a true decrease in measurable isotopic windows.

**Figure 5).**
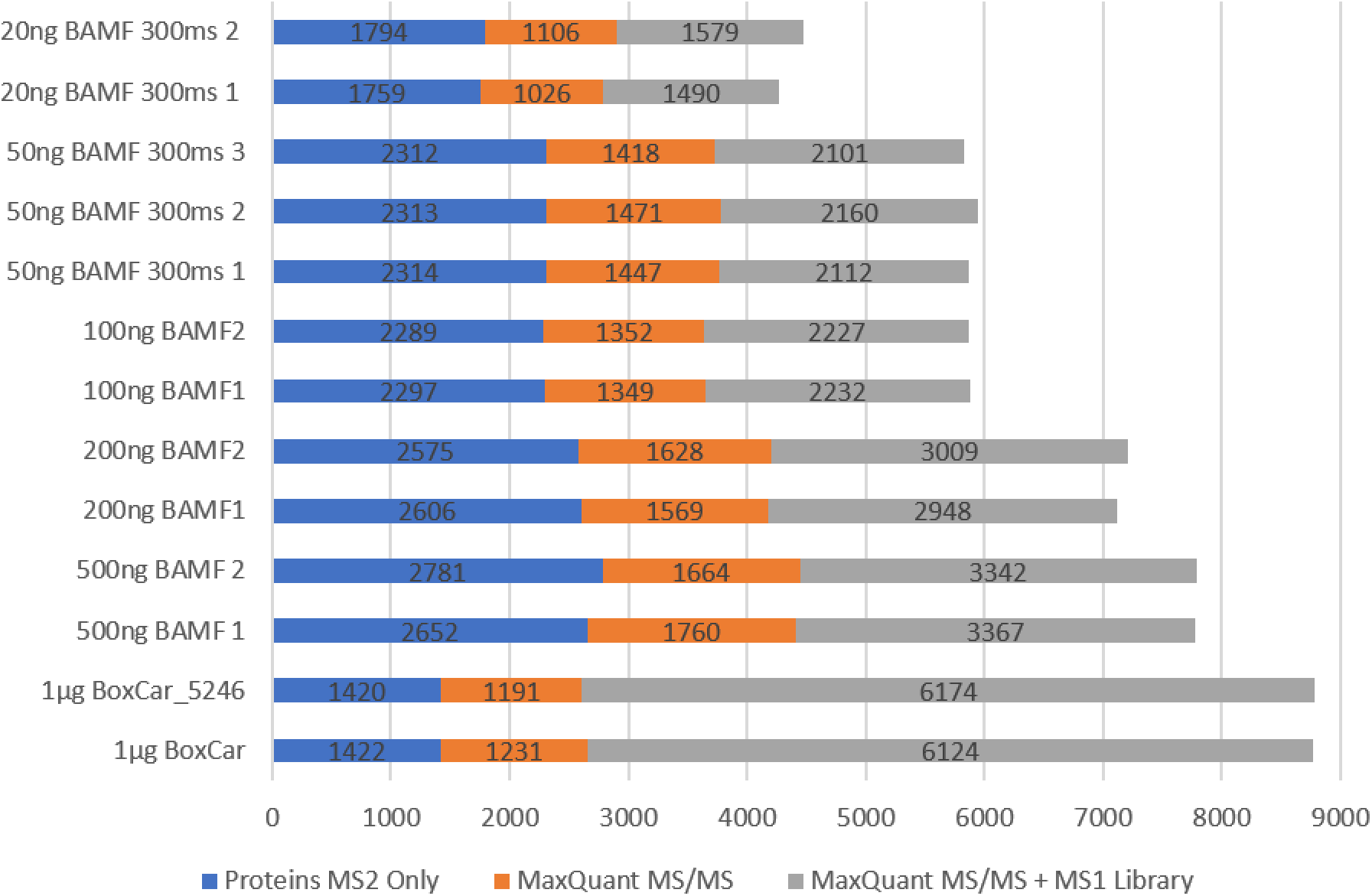
A composite of the number of proteins identified by Proteome Discoverer 2.2 in Blue, MaxQuant MS/MS FASTA matching in orange and with MS1 library matching in grey.

### Conclusions

We have described a method based on the BoxCar method that can be utilized on Orbitrap Tribrid systems with no modifications to the vendor software. Furthermore, we have expanded on the efficacy of the BoxCar method by directly picking ions for MS/MS from the BoxCar scans rather than from an additional MS1 full scan. The savings in time from the removal of the full scan coupled with the parallelization of MS1 and MS/MS acquisition using IRM HCD and ion trap acquisition allows for an increase in the number of MS/MS scans retrieved from each sample injection. Furthermore, we find that we can increase the dynamic range of peptides identified from both MS1 signal and from MS/MS fragmentation by using BAMF as compared to standard dd-MS2 experiments on the same Orbitrap system.

There are still limitations with the BoxCar and BAMF methods. Compared to standard dd-MS2, we obtain only a fraction of the number of MS/MS scans when peptide load on column is relatively high. However, for applications where only nanograms of material are available for use, BAMF may be an attractive alternative methodology. In our lab, work is underway to further develop BAMF for both metabolomics applications using an Orbitrap Fusion and Fusion ID-X systems as well as the newly released Exploris system that has integrated BoxCar/BAMF capabilities. The primary limiting factor is, however, informatics as improvements in the software for label free quantification including the recognition of BAMF scans by MaxQuant may be necessary for these techniques to realize their true potential.

## Acknowledgements

We would like to thank Dr. Sudipto Das and Dr. Maura O’Neill of NCI-Frederick for helpful conversations toward the construction of this manuscript.

Supplemental 1 PDF link

Supplemental 2 PDF link

Supplemental 3 PDF link

## References

(1) Meier, F.; Geyer, P. E.; Virreira Winter, S.; Cox, J.; Mann, M. BoxCar Acquisition Method Enables Single-Shot Proteomics at a Depth of 10,000 Proteins in 100 Minutes. Nat. Methods 2018. https://doi.org/10.1038/s41592-018-0003-5.

(2) Wichmann, C.; Meier, F.; Winter, S. V.; Brunner, A.; Cox, J.; Mann, M. MaxQuant.Live Enables Global Targeting of More than 25,000 Peptides. bioRxiv 2018. https://doi.org/10.1101/443838.

(3) Schilling, B.; Rardin, M. J.; MacLean, B. X.; Zawadzka, A. M.; Frewen, B. E.; Cusack, M. P.; Sorensen, D. J.; Bereman, M. S.; Jing, E.; Wu, C. C.; et al. Platform-Independent and Label-Free Quantitation of Proteomic Data Using MS1 Extracted Ion Chromatograms in Skyline. Mol. Cell. Proteomics 2012. https://doi.org/10.1074/mcp.M112.017707.

(4) Higgs, R. E.; Butler, J. P.; Han, B.; Knierman, M. D. Quantitative Proteomics via High Resolution MS Quantification: Capabilities and Limitations. Int. J. Proteomics 2013. https://doi.org/10.1155/2013/674282.

(5) Henry, H.; Sobhi, H. R.; Scheibner, O.; Bromirski, M.; Nimkar, S. B.; Rochat, B. Comparison between a High-Resolution Single-Stage Orbitrap and a Triple Quadrupole Mass Spectrometer for Quantitative Analyses of Drugs. Rapid Commun. Mass Spectrom. 2012. https://doi.org/10.1002/rcm.6121.

(6) Levy, M. J.; Washburn, M. P.; Florens, L. Probing the Sensitivity of the Orbitrap Lumos Mass Spectrometer Using a Standard Reference Protein in a Complex Background. J. Proteome Res. 2018. https://doi.org/10.1021/acs.jproteome.8b00269.

(7) Brunner, A. M.; Lössl, P.; Liu, F.; Huguet, R.; Mullen, C.; Yamashita, M.; Zabrouskov, V.; Makarov, A.; Altelaar, A. F. M.; Heck, A. J. R. Benchmarking Multiple Fragmentation Methods on an Orbitrap Fusion for Top-down Phospho-Proteoform Characterization. Anal. Chem. 2015. https://doi.org/10.1021/acs.analchem.5b00162.

(8) Scheltema, R. A.; Hauschild, J.-P.; Lange, O.; Hornburg, D.; Denisov, E.; Damoc, E.; Kuehn, A.; Makarov, A.; Mann, M. The Q Exactive HF, a Benchtop Mass Spectrometer with a Pre-Filter, High-Performance Quadrupole and an Ultra-High-Field Orbitrap Analyzer. Mol. Cell. Proteomics 2014. https://doi.org/10.1074/mcp.M114.043489.

(9) Hebert, A. S.; Richards, A. L.; Bailey, D. J.; Ulbrich, A.; Coughlin, E. E.; Westphall, M. S.; Coon, J. J. The One Hour Yeast Proteome. Mol. Cell. Proteomics 2014. https://doi.org/10.1074/mcp.M113.034769.

(10) Nagaraj, N.; Wisniewski, J. R.; Geiger, T.; Cox, J.; Kircher, M.; Kelso, J.; Pääbo, S.; Mann, M. Deep Proteome and Transcriptome Mapping of a Human Cancer Cell Line. Mol. Syst. Biol. 2011. https://doi.org/10.1038/msb.2011.81.

(11) Cox, J.; Hein, M. Y.; Luber, C. A.; Paron, I.; Nagaraj, N.; Mann, M. Accurate Proteome-Wide Label-Free Quantification by Delayed Normalization and Maximal Peptide Ratio Extraction, Termed MaxLFQ. Mol. Cell. Proteomics 2014. https://doi.org/10.1074/mcp.M113.031591.

(12) Doblmann, J.; Dusberger, F.; Imre, R.; Hudecz, O.; Stanek, F.; Mechtler, K.; Dürnberger, G. ApQuant: Accurate Label-Free Quantification by Quality Filtering. J. Proteome Res. 2018. https://doi.org/10.1021/acs.jproteome.8b00113.

