## Supplemental 1 for "BoxCar Assisted MS Fragmentation (BAMF)"

**Supplemental Table 1. BAMF 3 Window T-SIM Isolation windows**

**Experiment 1**

>>>>>>>>>>>>> Mass List Table <<<<<<<<<<<<<<

CompoundName|	Formula| AdductPositive|	m/z|	z|

| | +H| 400| 1|

| | +H| 520| 1|

| | +H| 640| 1|

| | +H| 760| 1|

| | +H| 880| 1|

| | +H| 1000| 1|

| | +H| 1120| 1|

| | +H| 1240| 1|

| | +H| 1360| 1|

| | +H| 1500| 1|

**Experiment 2**

>>>>>>>>>>>>> Mass List Table <<<<<<<<<<<<<<

CompoundName|	Formula| AdductPositive|	m/z|	z|

| | +H| 440| 1|

| | +H| 560| 1|

| | +H| 680| 1|

| | +H| 800| 1|

| | +H| 920| 1|

| | +H| 1040| 1|

| | +H| 1160| 1|

| | +H| 1280| 1|

| | +H| 1420| 1|

| | +H| 1540| 1|

**Experiment 3**

>>>>>>>>>>>>> Mass List Table <<<<<<<<<<<<<<

CompoundName|	Formula| AdductPositive|	m/z|	z|

| | +H| 480| 1|

| | +H| 600| 1|

| | +H| 720| 1|

| | +H| 840| 1|

| | +H| 960| 1|

| | +H| 1080| 1|

| | +H| 1200| 1|

| | +H| 1320| 1|

| | +H| 1460| 1|

| | +H| 1580| 1|
