## Supplemental 3 for "BoxCar Assisted MS Fragmentation (BAMF)"

Supplemental Table 3. BAMF 1 Window Method

Orbitrap Fusion Method Summary

Creator: Destroyer-Fusion\Orbitrap Fusion Last Modified: 6/28/2018 6:57:49 PM by BCO-Destroyer4/Fusion

Global Settings

Use Ion Source Settings from Tune = False

Method Duration (min)= 87

Ion Source Type = NSI

Spray Voltage: Positive Ion (V) = 2700

Spray Voltage: Negative Ion (V) = 2500

Sweep Gas (Arb) = 0

Ion Transfer Tube Temp (°C) = 300

APPI Lamp = Not in use

Pressure Mode = Standard

Default Charge State = 2

Advanced Precursor Determination = False

Experiment 1

Start Time (min) = 0

End Time (min) = 87

Cycle Time (sec) = 4.5

Scan tSIM

MSn Level = 2

Multiplex Ions Enabled = True

Maximum number of multiplexed ions = 10

Isolation Mode = Quadrupole

Define MSX IDs = True

Isolation Window (m/z) = 100

Loop Time= 3

Loop Count= 20

Loop Control= 3

Detector Type = Orbitrap

Orbitrap Resolution = 120K

Mass Range = Normal

Scan Range (m/z) = 350-1000

Maximum Injection Time (ms) = 200

AGC Target = 1000000

Inject ions for all available parallelizable time = True

Microscans = 1

RF Lens (%) = 60

Use ETD Internal Calibration = False

DataType = Profile

Polarity = Positive

Source Fragmentation = False

Scan Description =

>>>>>>>>>>>>> Mass List Table <<<<<<<<<<<<<<

CompoundName|	Formula| AdductPositive|	m/z|	z|	MSX ID|

| | +H| 430| 1| 1|

| | +H| 530| 1| 1|

| | +H| 630| 1| 1|

| | +H| 730| 1| 1|

| | +H| 830| 1| 1|

| | +H| 930| 1| 1|

| | +H| 1030| 1| 1|

| | +H| 1130| 1| 1|

| | +H| 1230| 1| 1|

| | +H| 1330| 1| 1|

Filter MIPS

MIPS Mode = Peptide

Filter ChargeState

Include undetermined charge states = False

Include charge state(s) = 2-6

Include charge states 25 and higher = False

Filter DynamicExclusion

Exclude after n times = 1

Exclusion duration (s) = 90

Mass Tolerance = ppm

Mass tolerance low = 5

Mass tolerance high = 5

Exclude isotopes = True

Perform dependent scan on single charge state per precursor only = False

Data Dependent Properties

Data Dependent Mode= Cycle Time

Scan Event 1

Scan ddMSnScan

MSn Level = 2

Isolation Mode = Quadrupole

Isolation Offset = Off

Isolation Window = 2.5

Reported Mass = Offset Mass

Multi-notch Isolation = False

Scan Range Mode = Auto Normal

FirstMass = 100

Scan Priority= 1

ActivationType = HCD

Is Stepped Collision Energy On = False

Stepped Collision Energy (%) = 5

Collision Energy (%) = 35

Detector Type = IonTrap

Ion Trap Scan Rate = Rapid

Maximum Injection Time (ms) = 300

AGC Target = 10000

Inject ions for all available parallelizable time = True

Microscans = 1

Use ETD Internal Calibration = False

DataType = Centroid

Polarity = Positive

Source Fragmentation = False

Scan Description =
